# Seeing danger? Visual constraints and collision risk in the Bearded Vulture *Gypaetus barbatus*

**DOI:** 10.64898/2026.11.03.763928

**Authors:** Simon Potier, Minglei Ma, Constance Blary, Sandrine Berthillot, Jérôme Cavailhes, Virginie M. Dos Santos, Ilka Champly, Marie Heuret, Thierry Bouchet, Alejandro Llopis Dell, Jean-Marc Lassance, Marjorie A. Liénard

## Abstract

The Bearded Vulture, *Gypaetus barbatus* is highly vulnerable to collisions with human-made aerial infrastructures. Yet, its visual sensory system remains poorly understood, limiting the development of species-specific collision mitigation measures. To address this gap, we characterized its visual ecology across four complementary dimensions: visual fields, spatial resolution, contrast sensitivity and spectral sensitivity of cone visual rhodopsins. Bearded vultures have no blind area in the direction of travel during foraging flight, indicating that frontal visual-field limitations are unlikely to contribute to their risk of collisions. Their axial length predicts high spatial visual resolution, suggesting that insufficient spatial resolution is unlikely to be a primary cause of collisions. By contrast, Bearded Vultures display very low peak contrast sensitivity compared to other birds, which may limit their ability to detect objects when contrast with the background, or within the objects themselves, is low. Functional characterization of the four cone visual pigments indicated sensitivity spanning approximately 350-640 nm, with a potential chromatic-discrimination range from 400 to 602 nm. Together, these results identify that contrast sensitivity, rather than visual-field configuration or spatial resolution, is a key perceptual constraint to consider in collision mitigation of the Bearded Vulture. Visual markers should therefore maximize both internal contrast and brightness contrast against the surrounding environment. More broadly, our multidisciplinary approach highlights how birds differ substantially in how they perceive anthropogenic structures and illustrates the importance of incorporating species-specific sensory ecology into conservation efforts, especially in the design of collision-mitigation strategies.

## Introduction

The global expansion of human-made aerial structures represents a major risk factor, causing various effects on birds: habitat loss and fragmentation, disturbances leading to displacement, alteration of migratory routes, and collisions with infrastructure (Drewitt & Langston, 2008). Collisions are a recognised major source of bird mortality worldwide (Rosenberg et al., 2019, Loss et al., 2014), especially with glass or other reflective building surfaces (Machtans et al., 2013), wind turbines (Marques et al., 2014, Bernardino et al., 2018, Drewitt & Langston, 2008), power lines and aerial cables that intrude into the open airspace (Rioux et al. 2013; Dos Santos, 2026). For instance, the number of bird deaths due to collision with wind farms can range from 0 to 40 individuals per turbine per year (Marques et al. 2014) and is correlated to bird behaviour and habitat, as well as wind turbine number and size (Thaxter et al. 2017).

Diurnal raptors of the order Accipitriformes are among the bird groups most vulnerable to wind turbine collisions, exhibiting the highest predicted collision rates among avian orders, with an average of 0.073 collisions per turbine per year (Thaxter et al. 2017). For perspective, this rate would correspond to approximately two collisions per year at a 30-turbine wind farm, a size which is approximately the median of individual European wind farms contributing to Accipitriform collision data in that study. Such a collision risk may ultimately threaten the viability of raptor populations (Duriez et al. 2023). This matter is becoming increasingly pressing, as the number of wind energy facilities continues to rise globally (Katzner et al. 2019; Conkling et al. 2022). Bird species with high wing loading such as vultures, bustards, or eagles are also particularly prone to collisions with power lines (SAIC 2000, Rioux et al. 2013, Eccleston and Harness 2018, Dwyer et al. 2018).

Different components of vision may contribute to collision risk in different species, making it crucial to identify the sensory mechanisms underlying collisions to develop effective mitigation strategies and reduce the impact of anthropogenic structures. Accordingly, characterizing bird visual abilities has emerged as a valuable framework for understanding why particular species are vulnerable to collisions and for informing effective mitigation strategies (Martin, 2011; Martin, 2022; Martin and Banks, 2023; Potier et al 2026). Indeed, vision enables birds to gather detailed and instantaneous spatial information from multiple directions and at varying distances (Land & Nilsson, 2012; Martin, 2017). Raptors, such as the Bearded Vulture, *Gypaetus barbatus*, are believed to be highly visually dependent (Jones et al. 2007, Mitkus et al. 2018, Potier et al. 2020). Among all animals studied so far, Old-World vultures and the Wedge-tailed eagle *Aquila audax*, for instance, possess the highest known spatial visual resolution, *i.e.* the ability to resolve fine details in the visual field (Fischer 1968, Reymond 1985), but it is now known that distinct raptors differ highly in their spatial resolution of vision (Martin 2017, Caves et al. 2024). Furthermore, broad variation in visual abilities such as contrast sensitivity (Blary et al. 2024) and visual fields have been identified (Potier et al. 2018a). In particular, while the Egyptian vulture *Neophron percnopterus* is able to see directly ahead during flight (Potier et al. 2018a), most *Gyps* vultures possess a frontal visual blind zone that extends into their flight path during foraging (Martin et al. 2012). As a result, they are unlikely to detect obstacles located directly ahead, potentially increasing their risk of collision with structures encountered during flight. This substantial variation in raptors’ visual abilities (reviewed in Potier et al. 2020) may help explain interspecific differences in susceptibility to collisions.

The Bearded Vulture is a large, long-lived Accipitriform scavenging raptor inhabiting mountainous regions of Eurasia and Africa (Figure 1) and is highly specialized for an osteophagous diet consisting predominantly of bones from ungulate carcasses. It belongs to the Gypaetinae, a distinct evolutionary lineage of Old-World Vultures that differs markedly from the more typical Aegypiinae vultures in its ecology and behaviour (Wink et al 2026). The species is currently classified as Near Threatened on the IUCN Red List and has a highly fragmented distribution in Europe. The European population is currently estimated at 617 individuals, including between 316 to 419 individuals in the Alps in 2024 (Lauper 2024). Collisions account for 18% of the species’ mortality in Europe (Margalida et al. 2008). This threat is particularly concerning because an increase in mortality of only four adult females in the Alpine population would lead to a population decline in the absence of supplementary reintroductions, captive-bred releases, or the translocation of individuals from more abundant wild populations (Schaub et al. 2024). Bearded Vultures perform approximately 74% of their flight trajectories below 200 m above ground level, an altitude range at which birds are more likely to encounter human-made structures, such as cables or wind turbines, increasing the risk of collisions (Vignali et al. 2022; Figure 1).

**Figure 1.**
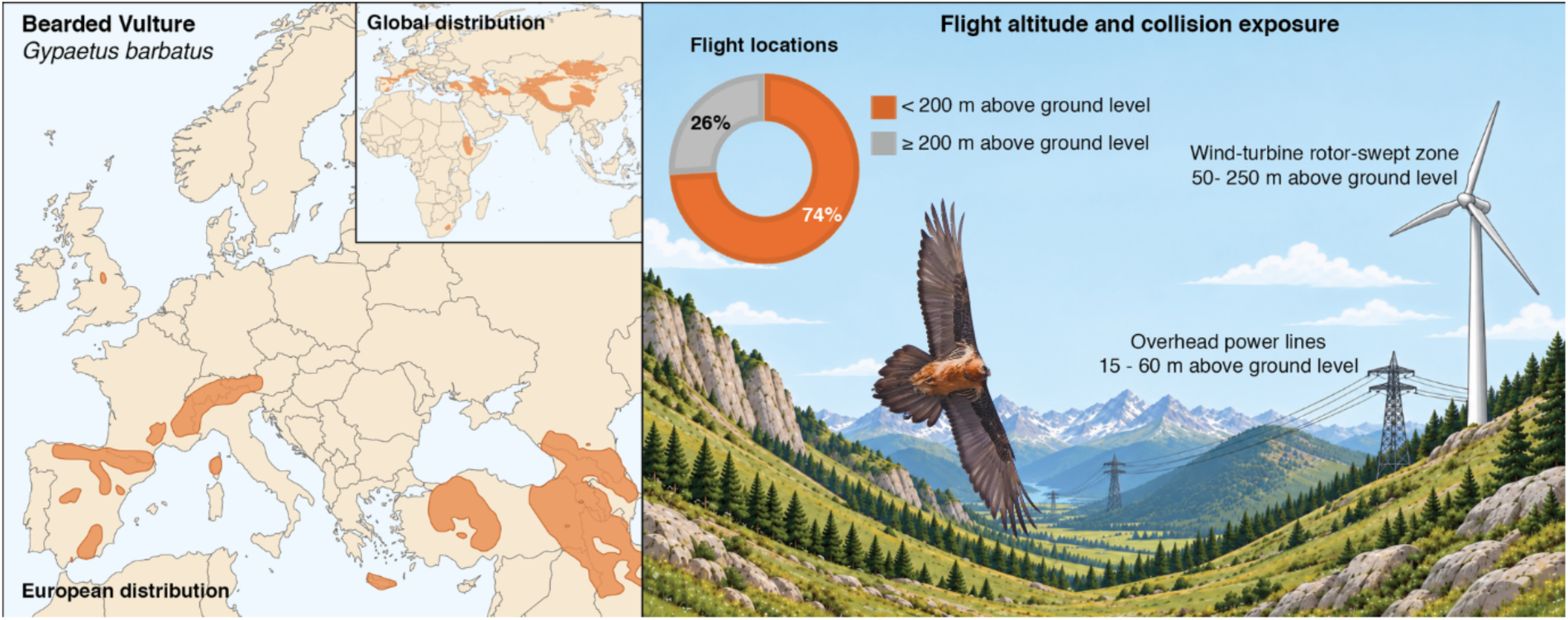
Distribution, flight altitude and potential collision exposure of the Bearded Vulture. (*left panel*) The European distribution of the Bearded Vulture is shown together with its global distribution (inset). Distribution data correspond to the 2023 resident range from eBird Status and Trend (Fink et al. 2024). (*right panel*) Schematic illustration of the relationship between Bearded Vulture flight altitude and potential exposure to collision hazards from anthropogenic aerial infrastructures. The landscape and infrastructure are ai-generated. Bearded Vulture photograph courtesy of Mr Wildclicks, via Canva Pro. Flight-altitude frequencies from Vignali et al. (2022).

To better understand the visual mechanisms that may contribute to collision risk in the Bearded Vulture, we examined its visual fields, visual acuity, contrast sensitivity, and the wavelength of absorbance of its cone visual pigments. Characterizing these components of the visual system provide important insights into why birds collide with aerial structures (Martin 2022), to suggest concrete mitigation adapted to their ecology and behaviour, as recently shown in the Black grouse *Lyrurus tetrix* (Potier et al. 2026) and the Red kite *Milvus milvus* (Potier et al. in revision).

## Material and methods

### Birds and study locations

Experiments were conducted on three Bearded Vultures in accordance with the European Union Directive (2010/63/EU) on the protection of animals used for scientific purposes. The experimental protocol was approved by the Languedoc-Roussillon Ethics Committee no. 36 under reference 2148-00000. Measurements were conducted in February 2024 at Puy du Fou France (France) on an immature female less than one year old, and in October 2024 at the Centre de Fauna Vallcalent (Lleida, Spain) on an adult male and an adult female. All measurements were taken on site, after which the birds were returned to their aviaries. These three individuals were the only birds made available for the experiments to minimize disturbance on the ongoing captive breeding programs.

### Visual field

The visual field characteristics of each bird were evaluated using the ophthalmoscopic reflex technique, following established protocols (Martin and Piersma 2009). Briefly, each bird was restrained for 20-30 min in a plastic tube of the appropriate size to minimize body movement and the bill was placed in a custom-made bill holder made of steel and aluminium. The head was positioned to approximate the posture spontaneously adopted by the birds when held in the hand. The surfaces of the bill holder were coated with cured silicone sealant to provide a non-slip, cushioned surface. The bill was held in place with Micropore^TM^ tape, leaving the nostrils unobstructed to allow normal breathing. Calibrated dorsal and lateral photographs of the head and eyes were taken using an Iphone 13 (Apple Inc., CA, USA) to determine eye position within the skull (Martin and Piersma 2009).

The head was held in position at the centre of a visual perimeter - a device that permits the eyes to be examined from known angular positions relative to the head. The perimeter coordinate system followed conventional latitude and longitude measures, with the equator oriented vertically in the median sagittal plane of the head, which divides the head symmetrically into left and right hemispheres. This coordinate system was subsequently used to present the visual field data. The eyes were examined using an ophthalmoscope mounted on the perimeter arm, allowing measurements with an accuracy of ± 0.5°. The boundaries of the retina projection were measured using the positions that the eyes spontaneously adopted when they were rotated fully forwards (converged) to estimate the binocular area boundaries and backwards (diverged) to estimate the blind sector directly behind the head. We did not measure the degree of eye movements or the pecten projection to minimize restraint time.

A correction was applied to the measurements considering a hypothetical viewing point placed at infinity following Martin (1984). This correction accounts for the distance between the head and the visual perimeter used in the measurements taken with the visual field apparatus and the horizontal separation between the eyes (Martin, 1984). A topographical map of the visual field and its different components was then derived, including lateral fields, binocular field and blind areas above and behind the head. The limits of the visual field were determined at 10° intervals of elevation in an arc, enabling a complete estimation of the binocular field sector. At elevations where the bill holder obstructed the view, the binocular field width was estimated as the mean value of the measurements immediately above and below the obstructed elevation (Martin, 2007).

### Axial length and spatial resolution

Across birds, axial length is a strong predictor of spatial resolution (Kiltie 2000; Caves et al 2024). Axial length (AL) in the Bearded Vulture has been estimated at 30.3 mm (Ritland, 1982). Spatial resolution (SR; cycles degree^-1^) was then estimated from AL (mm) using the relationship for diurnal species reported by Kiltie (2000):

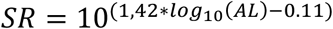

It should be noted that a more recent relationship between axial length and spatial resolution has been derived from a large avian dataset while accounting for phylogeny (Caves et al. 2024). However, the applicability of this general model to raptors remains uncertain, with substantial deviations between predicted and empirically measured acuity for several species (Figure S1). Both the Caves et al. (2024) and Kiltie (2000) relationships tend to underestimate measured spatial resolution in Accipitriformes vultures for which empirical acuity estimates are available (Egyptian Vulture *Neophron percnopterus* and Griffon Vulture *Gyps fulvus*), but the Kiltie relationship consistently provides estimates closer to the observed values (Figure S1). Given that the Kiltie relationship more closely predicts empirical spatial resolution in these Accipitriformes vultures, together with the close phylogenetic relationship between the Bearded Vulture and the Egyptian Vulture (Wink et al. 2026), we therefore considered the Kiltie (2000) relationship more appropriate for estimating the spatial resolving power of the Bearded Vulture.

### Contrast sensitivity

We used a reflex-like behaviour setup to study the contrast sensitivity of each individual, following Wagner et al. (2022) and Blary et al. (2024). The experimental set up, stimuli and experimental procedures are detailed in Supplementary methods.

### Opsin functional characterisation in HEK293T cells

Opsin protein sequences were predicted from the publicly available Bearded Vulture genome assembly bGypBar2.pri (NCBI accession GCA_028022735.1) as detailed in Supplementary methods. Predicted amino acid sequences of the four visual opsins uncovered are provided in supplementary material. The visual cone opsins (SWS1, SWS2, Rh2, LW) open reading frames were codon-optimized and subcloned into pcDNA5-FLAG-T2A-mRuby2 (Liénard et al., 2021). Prior to obtaining each rhodopsin dark spectrum, opsin expression and localisation was verified via immunofluorescence as detailed in supplementary methods. Rhodopsin complexes were then reconstituted in human embryonic kidney cells (HEK293T) in presence of 11-*cis*-retinal following the PASHE procedure (Liénard et al., 2022; Roberts et al., 2025) as detailed in supplementary methods.

## Results

A summary of the visual parameters measured or estimated for the Bearded Vulture is presented in Table 1.

**Table 1.** Summary of measured and estimated visual parameters of the Bearded Vulture (*Gypaetus barbatus*).

| **Visual parameter** | **Value** |
|---|---|
| ***Measured visual fields (Figure 2)*** |  |
| Horizontal field at rest | 313 ± 3.9° |
| Lateral fields | 143 ± 5.8° |
| Frontal binocular overlap | 27 ± 2.1° |
| Maximal binocular overlap at 30° above bill-tip | 34 ± 3.0° |
| Blind horizontal area behind head | 47 ± 3.9° |
| Vertical binocular field | 135°<sup>a</sup> |
| Binocular field width above the head (at 0°) | 6.5 ± 9.5°<sup>b</sup> |
| ***Estimated spatial Resolution (SR)*** |  |
| Axial length (mm) | 30.3 |
| Estimated spatial resolution in bright daylight (cycles per degree) | 98.5 |
| Estimated spatial resolution in crepuscular conditions (cycles per degree; reduced 10-fold) | 9.85 |
| ***Measured contrast sensitivity (Figure 3)*** |  |
| Peak contrast sensitivity (1/Cm) | 9.5 |
| Michelson contrast threshold at peak sensitivity (Cm) | 0.105 (10.5%) |
| Spatial frequency at peak contrast detection limit (cycles per degree) | 1.27 |
| ***Measured visual pigment maximum absorbance (λ<sub>max</sub>) (Figures 4 and 5)*** |  |
| SWS1 | 401 nm |
| SWS2 | 446 nm |
| Rh2 | 505 nm |
| LW | 542 nm |
| Upper sensitivity overlaps between Rh2 and LW (colour vision limit) | 602 nm |
| Maximal upper wavelength perception | 639 nm |

<sup>a</sup> This value should be considered an approximation, as the experimental apparatus does not allow precise measurement of the vertical extent.
<sup>b</sup> This value should be interpreted with caution given the marked inter-individual variability (see Figure 2).

### Visual field

The maximum binocular overlap of the Bearded vulture was 34° and occurred 30° above the bill-tip direction (Figure 2A). In the horizontal plane, the visual field covered 313°, comprising a frontal binocular overlap of 27°, lateral fields of 143° on each side, and a blind sector of 47° behind the head (Figure 2B). In the median sagittal plane, the binocular field extended vertically from 0° to 105° in between the head tip (0°) and the bill-tip (100°), 10° behind the head tip, and 20° below the bill-tip plane (Figure 2C). Thus, the binocular field provided a view of 135° vertically, encompassing the path forward while also providing binocular vision above the head and beyond the bill-tip position during flight (Figure 2D).

**Figure 2.**
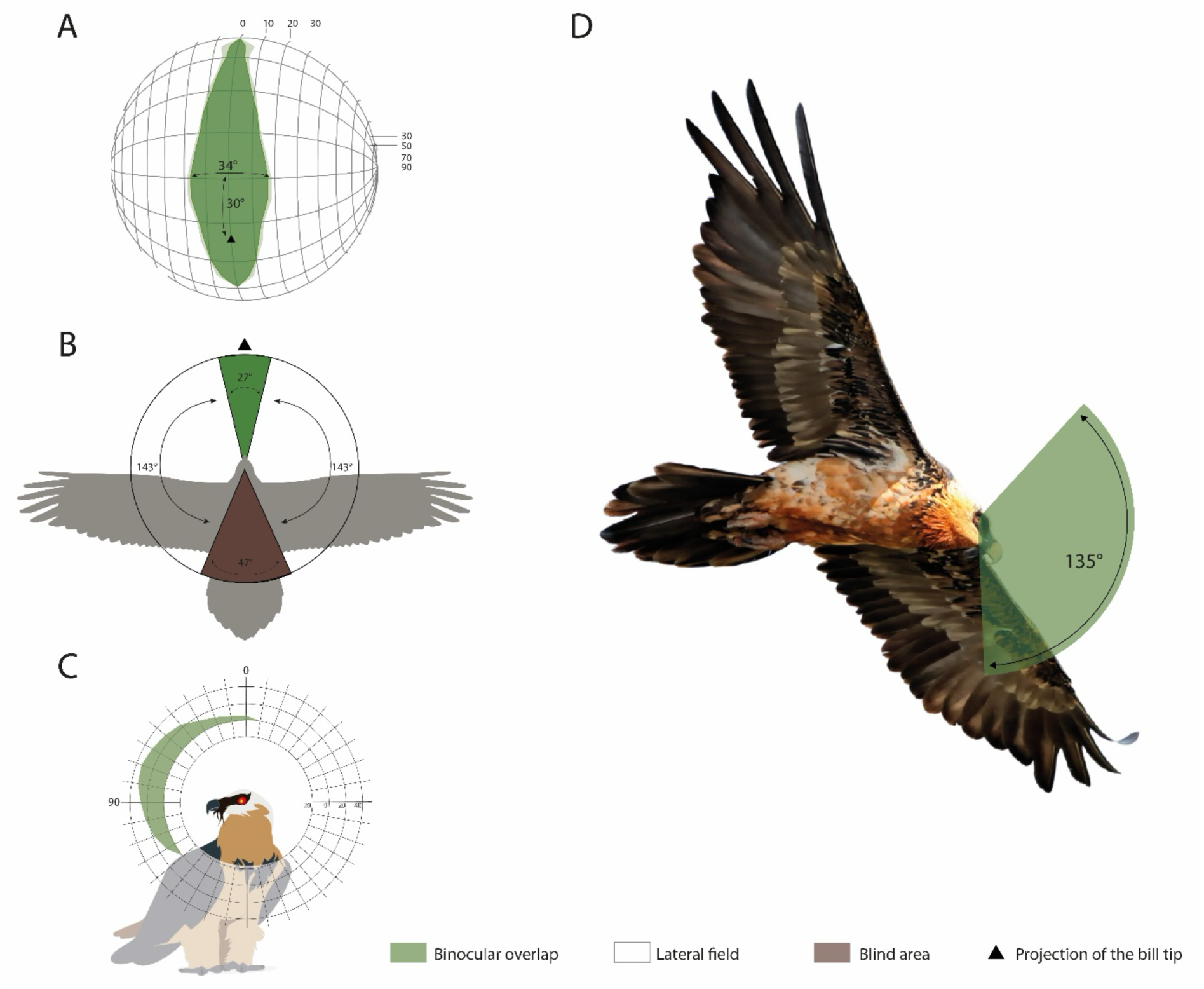
Visual fields of the Bearded Vulture. (**A**) Orthographic projection of the retinal field boundaries of both eyes. A latitude and longitude coordinate system is used, with the equator aligned vertically in the median sagittal plane. The bird’s head is positioned at the centre of a schematic globe, and the projection of the bill tip corresponds to the black triangle. Grid lines are drawn at 10° intervals in latitude and at 20° intervals in longitude. Dark green areas represent the binocular sector; light green areas correspond to the standard deviation. (**B**) Section through the horizontal plane corresponding to the bold line on panel A, showing the binocular overlap (green), lateral fields (white) and blind area (brown) in Bearded Vulture. (**C**) Binocular field as a function of vertical elevation in the median sagittal plane. The bird’s head orientation is indicated by the direction of the bill tip. All values are in degrees, and the grid is at 10° intervals. Values represent the mean visual field of the three measured individuals. **(D)** Photograph of a Bearded vulture in foraging flight showing the typical angle of the eye-bill tip projection when foraging. Images by Wirestock on Freepik and EcoPic on Getty Images. Silhouette by PhyloPic.org.

### Axial length and spatial resolution

Based on the relationship established by Kiltie (2000) we predicted a spatial resolution (SR) for the Bearded Vulture at 98.5 cycles/degree, while recognizing the relative uncertainty associated with morphology-based predictions (Figure S1).

### Contrast sensitivity

To assess the bird’s ability to distinguish features in its visual environment, we projected stripe patterns with varying contrast and spatial frequencies. Comprehensive data was obtained for the male individual, whereas the two female individuals showed weaker responses to the visual stimuli. Partial measurements were obtained for the young female originating from Vallcalent, while no contrast-sensitivity data could be obtained for the adult female from France.

Peak contrast sensitivity of the Bearded vulture was 9.5 at a spatial frequency of 1.27 cycles/degree (Figure 3A). This sensitivity corresponds to a Michelson contrast (C_m_) threshold of 0.105 (10.5%), which means that Bearded Vulture cannot distinguish an object from the surrounding background unless it is at least 11% brighter. Compared to other Accipitriformes for which equivalent measurements are available, the Bearded Vulture exhibited the lowest peak contrast sensitivity reported to date (Figure 3B).

**Figure 3.**
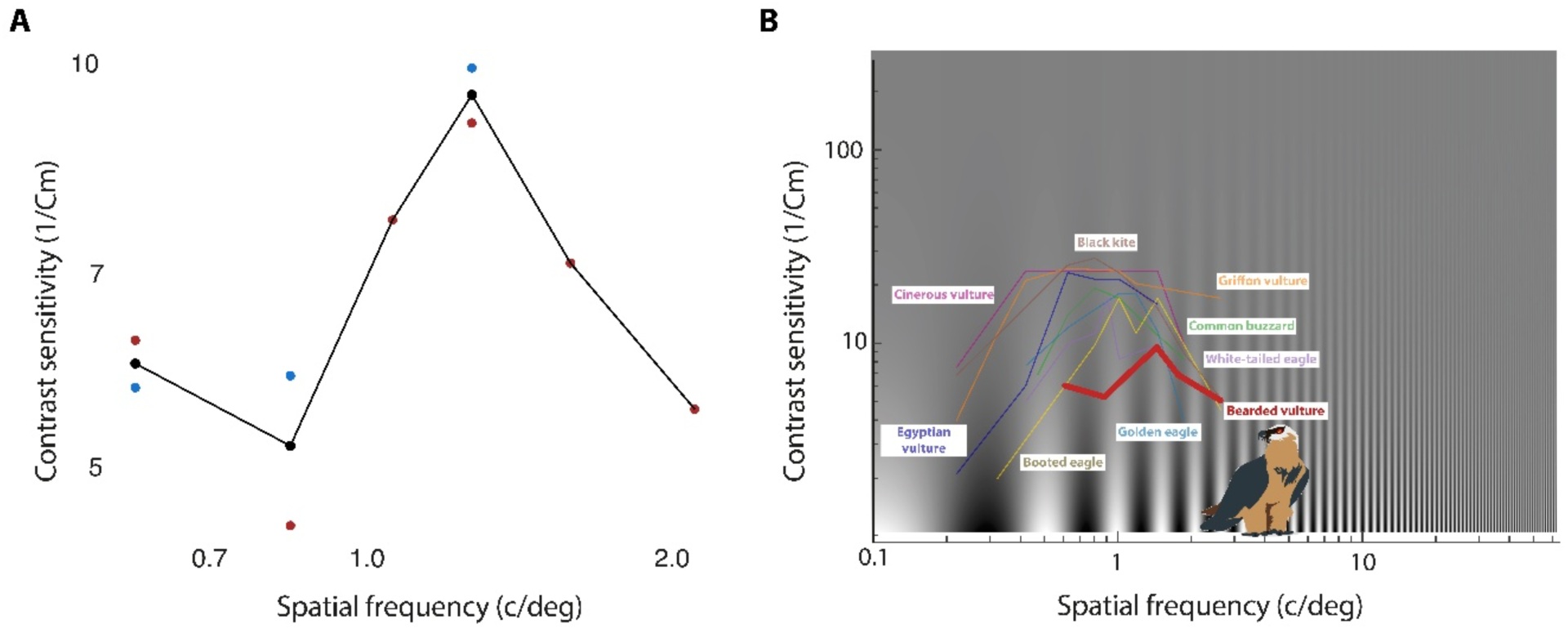
Contrast sensitivity of the Bearded Vulture. Contrast sensitivity, defined as the inverse of contrast threshold, is a function of spatial frequency. (**A**) Red and blue points represent contrast sensitivity for the male and female individual measured, respectively, and were calculated as the inverse of Michelson Contrast (1/C_m_) values. The black curve and points represent the mean of the available individual measurements at each spatial frequency. (**B**) Comparison of the contrast sensitivity function of the Bearded Vulture (red curve) with those of other Accipitriformes using data from Blary et al. (2024): Egyptian Vulture *Neophron percnopterus*, Cinereous Vulture *Aegypius monachus*, Griffon Vulture *Gyps fulvus*, Black Kite *Milvus migrans*, White-tailed Eagle *Haliaeetus albicilla*, Golden Eagle *Aquila chrysaetos*, Booted Eagle *Hieraaetus pennatus* and Common Buzzard *Buteo buteo*.

### Spectral absorbance of cone visual rhodopsins and estimated spectral sensitivity

We characterized the spectral absorbance of the four Bearded Vulture cone visual pigments (SWS1, SWS2, Rh2, LW). Opsins expressed in HEK293T cells and reconstituted in presence of 11, *cis*-retinal formed functional visual pigments with estimated wavelengths of maximal absorbance (λ_max_) of 400.8 nm (95% CI: 399.7-401.8 nm) for SWS1 (Figure 4A), 446.3 nm (443.9-448.6 nm) for SWB2 (Figure 4B), 504.6 nm (496.9-511.2 nm) for Rh2 (Figure 4C), and 541.6 nm (537.4-545.7 nm) LW (Figure 4D, Figures S2-S3)

**Figure 4.**
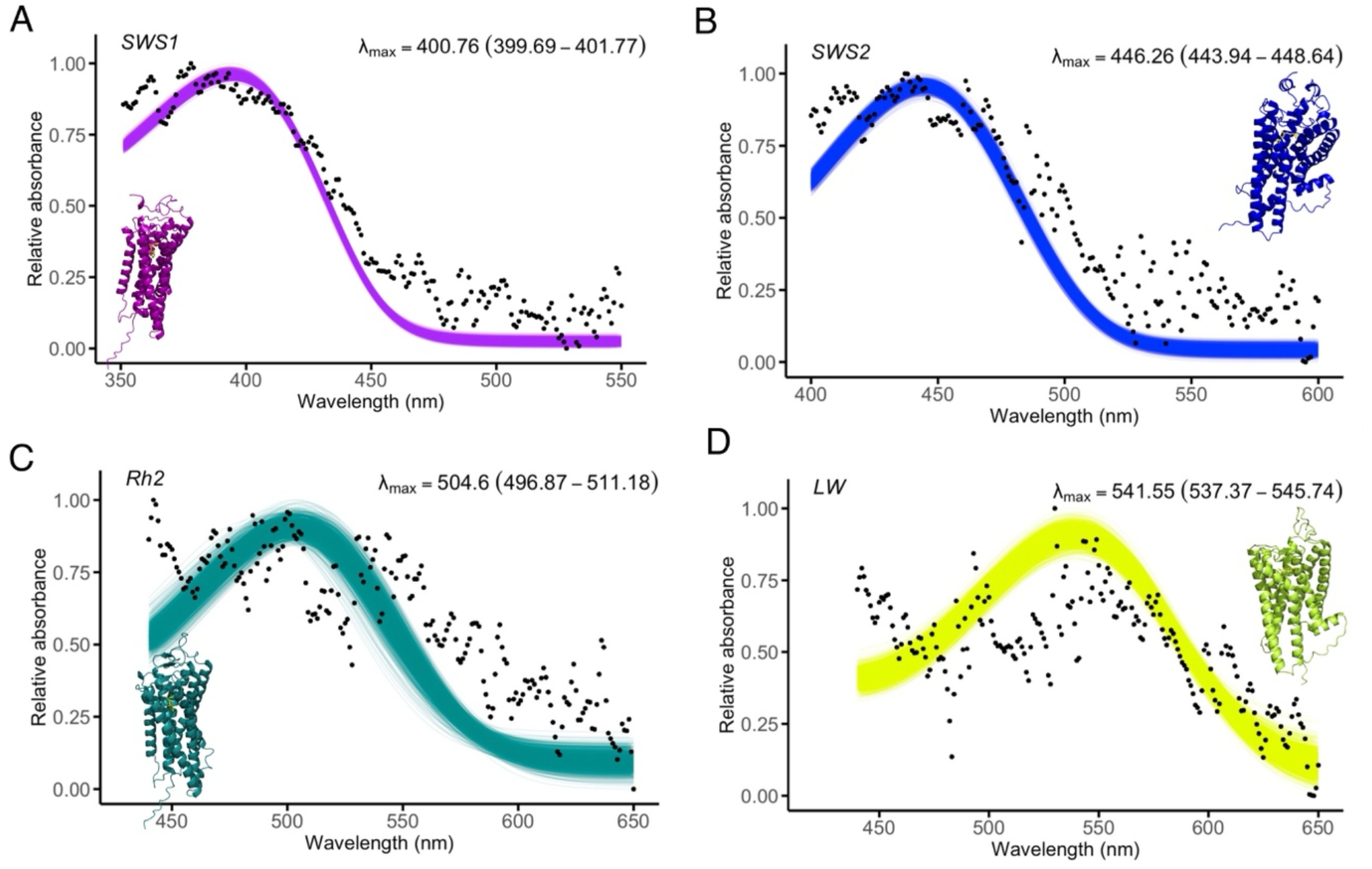
Functional expression and spectral characterization of Bearded Vulture cone visual pigments. UV-Vis absorbance spectra of the reconstituted and purified short-wavelength-sensitive (SWS1 and SWS2), middle-wavelength-sensitive (Rh2), and long-wavelengh-sensitive (LW) visual pigments expressed in HEK293T cells. Visual pigments were reconstituted with 11-*cis*-retinal and purified under dark conditions (see also Figs S2 and S3). Black dots represent mean relative absorbance at each wavelength across replicate measurements for **(A)** SWS1 (*n* = 6), **(B)** SWS2 (*n* = 6), **(C)** Rh2 (*n* = 2), and **(D)** LW (*n*=7), where *n* is the number of measurements of protein aliquots containing rhodopsins. Relative absorbance spectra were fitted using the Govardskii visual-pigment template. Coloured curves represent the fits obtained from 1,000 bootstrap replicates, with colours corresponding approximately to the human-perceived colour associated with the estimated λ_max_ of each pigment. Estimated λ_max_ and their confidence intervals are indicated in each panel. Opsin membrane localisation and protein purification fractions are presented in Figures S2 and S3, respectively.

The fitted λ_max_ values were subsequently used to reconstruct theoretical relative absorbance spectra using the Govardovskii visual-pigment template (Figure 5). For each visual pigment, the upper wavelength sensitivity limit was then theoretically predicted by applying a 5% sensitivity cut-off to the upper absorbance shoulder of the opsin. This yielded an upper limit of long-wavelength sensitivity of 639 nm for the visual system. Using the corresponding 5% threshold of Rh2 as a theoretical limit for potential chromatic-discrimination between Rh2 and LW cones yielded an upper wavelength of 602 nm, corresponding approximately to the orange-amber region of the human-visible spectrum. At shorter wavelengths, the theoretical lower boundaries of potential cone detection and chromatic discrimination were estimated at approximately 350 and 400 nm.

**Figure 5.**
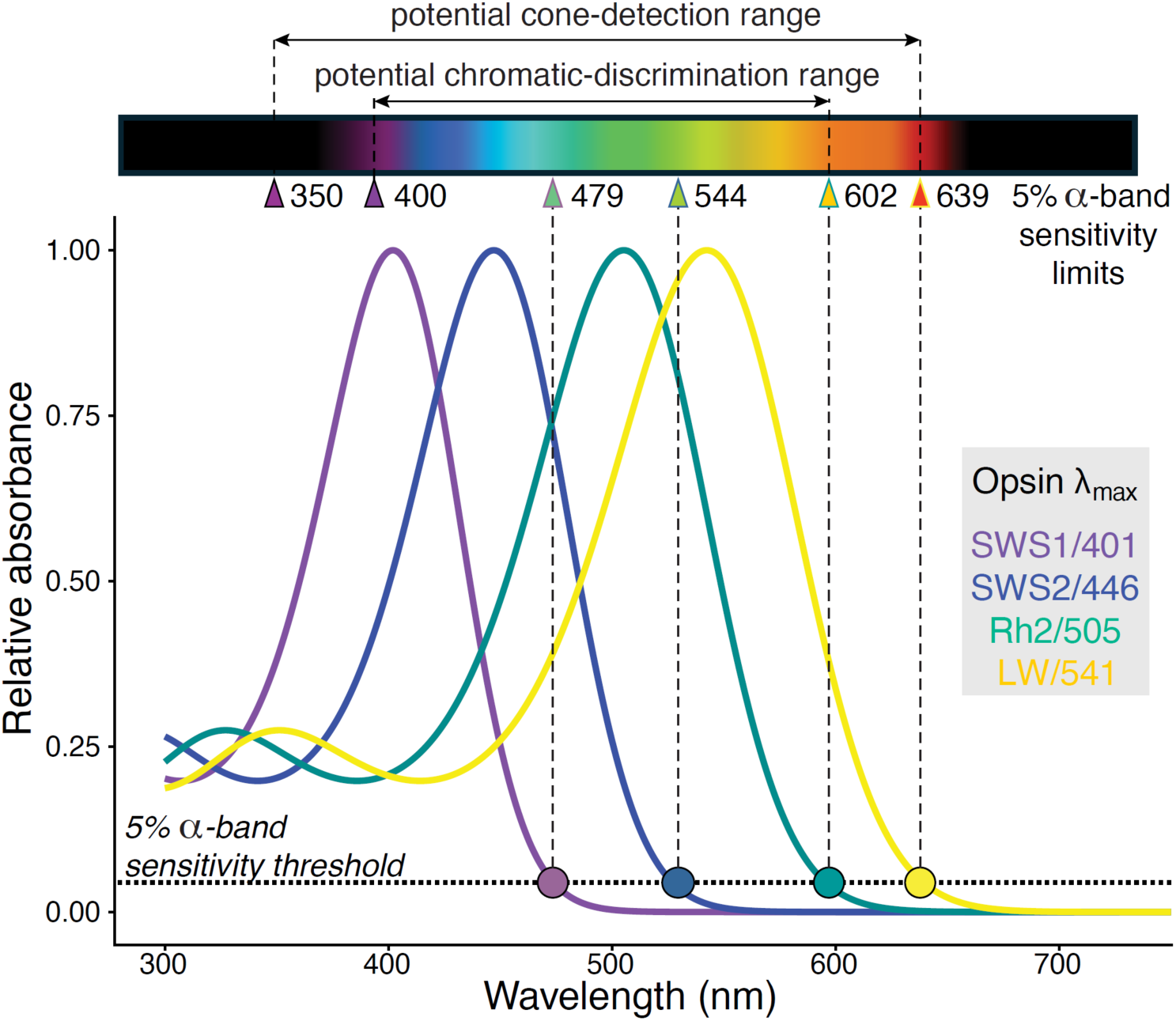
Spectral sensitivity of Bearded Vulture based on cone visual pigments. Theoretical relative absorbance spectra were reconstructed from the fitted *λ*_max_ values of SWS1, SWS2, Rh2 and LW using the Govardovskii template for visual pigments. The 5% points of the *α*-band were used as operational criterion to estimate the limits of cone spectral sensitivity, excluding secondary *β*-band absorbance at short wavelengths. Coloured circles indicate the 5% sensitivity limits on the long-wavelength portion of the spectrum of each pigment, while arrowheads indicate these limits as well as the corresponding short-wavelength limits of SWS1 and SWS2. The potential cone-detection range was defined by the 5% *α*-band sensitivity limits of the outermost cone pigments, extending from approximately 350 nm for SWS1 (estimated from the relative 5% absorption level of its ascending curve, corresponding to ∼0.25 absorbance) to 639 nm for LW. The potential chromatic-discrimination range was conservatively defined as the wavelength interval over which at least two cone pigments retain >5% *α*-band sensitivity, extending from approximately 400 nm, the short-wavelength limit of SWS2, to 602 nm, the long-wavelength limit of Rh2. These ranges represent theoretical limits based on visual-pigment absorbance and do not account for filtering in the ocular media or oil droplets, receptor noise, or post-receptor processing.

## Discussion

To gain a more detailed understanding of the abilities and constraints imposed by the Bearded Vulture visual system, we quantified its visual binocular field, corneal diameter as a morphological predictor of spatial resolution, contrast sensitivity, and the spectral sensitivity of its cone visual pigments. Together, these complementary components of vision provide insight into the sensory mechanisms that may contribute to collision risk and inform the design of mitigation measures that account for the visual capabilities of the Bearded Vultures

In the context of avian collisions both the binocular overlap, which provides information about the direction of travel, and the vertical extend of the binocular field, which determines vision directly along the flight path, are important to consider. The Bearded Vulture frontal binocular field covers 27° in the horizontal plane, falling within the average range observed in birds (20–40°; Martin, 2017). This value is also typical amongst most scavenger species (Potier et al. 2018a), except some *Gyps* vultures, which have a narrower binocular overlap (Martin et al. 2012), potentially reflecting differences in foraging ecology between solitary, specialized foragers such as the Bearded Vultures and colonial *Gyps* vultures (Martin, 2017; Oliva-Vidal et al 2024). Binocular vision may also support precise bill placement during feeding (Martin, 2009). The maximum binocular overlap, which reaches 34° and is positioned 30° above the bill-tip direction, is similar to most raptors that forage in flight (Potier et al., 2018a). The vertical extent of the binocular field (135°) is particularly notable for a raptor, with only the Turkey Vulture *Cathartes aura* showing a comparable specialization (Potier et al., 2018a). Therefore, in contrast to other collision-prone species sensitive such as *Gyps* vultures (Martin et al. 2012), Red Kites *Milvus milvus* (Potier et al. in revision) or mature males Capercaillies *Tetrao urogallus* (Potier et al. in prep), Bearded Vultures do not experience a blind area in the direction of travel when foraging (Figure 2). This distinction is important for collision mitigation because it suggests that obstacles and visual markers positioned within the forward flight path are potentially visible to Bearded Vulture, provided that they are sufficiently conspicuous.

The predicted spatial resolution of the Bearded vulture is among the highest reported to date (98.5 cycles/degree), closely approaching the range of other vultures such as the Indian Vulture *Gyps indicus* (135 cycles/degree), the Griffon Vulture *Gyps fulvus* (104 cycles/degree), and the Egyptian Vulture *Neophron percnopterus* (135 cycles/degree) (Fischer 1968). For comparison, the more recent general avian relationship proposed by Caves et al. (2024) - which show substantial deviations between predicted and empirically measured in accipitriforms - yielded a substantially lower estimate of 28.8 cycles/degree, well below the values reported for other Accipitriformes vultures. We therefore retained 98.5 cycles/degree as a primary estimate of Bearded Vulture spatial resolution. Such high spatial resolution is consistent with the visual demands of searching for food while soaring, with Bearded Vultures foraging at altitudes comparable to those of Griffon Vultures in the Alps (Genero 2017, Vignali et al. 2022). Although direct measurement will ultimately be required to determine Bearded Vulture visual acuity, its large eyes are consistent with relatively high spatial resolving power. Limited spatial resolution, therefore, seems unlikely to represent a major visual constraint contributing to collision risk in this species.

By contrast, the Bearded Vulture exhibits a notably low peak contrast sensitivity (9.5) compared with other raptors, including the Cinereous Vulture (*Aegypius monachus*: 25.6), Griffon Vulture (22.6), and the Egyptian Vulture (20.4) (Blary et al. 2024). Peak contrast sensitivity in birds is typically lower than that of mammals (e.g., approximately 200 in humans; De Valois et al. (1974)) and ranges from 4.9 in the Chilean Flamingo *Phoenicopterus chilensis* to 31 in the American Kestrel *Falco sparverius* (Hirsch 1982, Blary et al. 2024). Thus, with a peak contrast sensitivity approximately 20-fold lower than that of humans, Bearded Vultures may have difficulty detecting aerial objects when their contrast against the visual background is low. This limitation is also strongly dependent on viewing conditions, including illumination, atmospheric conditions, viewing distance, and the visual background. For instance, unlike humans, Black Grouse *Lyrurus tetrix* cannot perceive some of the internal contrasts of visual markers placed on ski lift cables (Potier et al. 2026). Therefore, improving both the internal achromatic contrast of aerial human-made objects (such as wind turbines), and visual markers in particular, as well as their contrast against the surrounding environment may be particularly important for reducing collision risk, as also supported in previous studies (McIsaac 2001, Hodos 2002, May et al. 2020, Martin 2022). Indeed, increasing internal contrast and ensuring robust contrast against the environment has been shown to enhance bird detectability of rotating wind turbine blades (Blary et al. 2023, Blary et al. 2026).

The spectral characteristics of the Bearded Vulture visual pigments align with avian pigment sensitivities (reviewed in Hagen et al. (2023)), and together its four avian cone opsins absorb wavelengths of light spanning ultraviolet to red from ca. 350 to 639 nm. Although ocular media transmittance could not be measured on intact eyes directly, some UV light is expected to reach the retina, as is the case in most bird species possessing a near-UV-sensitive SWS1 visual pigment sensitivity (reviewed in Hart and Voroboyev 2005). Consequently, the Bearded Vulture eye is likely to perceive at least some luminance (brightness) of a narrow range of ultraviolet-violet wavelengths invisible to the human eye. This capability may be particularly relevant because achromatic (luminance-based) cues can be seen over greater distance than chromatic cues (Lind & Kelber, 2011, Potier et al., 2018b). Therefore, incorporating UV-reflective components into collision mitigation devices could enhance their long-range detectability while largely remaining imperceptible to humans, effectively providing a specific channel for obstacle detection during flight. Chromatic contrasts and spectral sensitivity are still important to consider in collision context (Fernández-Juricic et al., 2011, Zheng et al., 2022). For practical reasons, absorbance measurements of Y and R oil droplets in Rh2 and LW cones from fresh eyes in this endangered species were not possible during this study. The long-wavelength limit of cone sensitivity is largely unaffected by oil-droplet filtering (Hart and Vorobyev 2005). Together with the descending limb of absorbance of the expressed Rh2 cone rhodopsin intersecting the long-wavelength absorbance curve, this places the theoretical limit of colour vision in the Bearded Vulture at approximately 602 nm, where it would perceive wavelength of yellow or orange bird markers robustly. Colour vision requires opponency between photoreceptors with distinct, overlapping spectral sensitivities. Long-wavelength colour discrimination may therefore extend into the deep-orange range, potentially allowing discrimination of yellow or orange markers. This contrasts with the more limited long-wavelength sensitivity predicted for some ground-nesting species, such as mountain Galliformes (Potier et al 2026).

Regarding achromatic vision, although the Bearded Vulture, is unlikely to perceive deep red wavelengths with the same sensitivity as humans, its luminance-sensitive photoreceptors could retain sensitivity up to ca. 640 nm. This may provide the eye with detectable contrast information from red or orange markers placed on aerial cables with the surrounding background *i.e* the sky seen under different visibility conditions. This long-wavelength sensitivity is consistent with the behavioural relevance of ferruginous colouration in this species, as Bearded Vultures actively seek out iron oxide-rich substrates to acquire their characteristic red-orange plumage and show a preference for orange-red substrates when offered different colours (Negro et al 1999; Wink et al 2026). However, whether these substrates provide chromatic or luminance cues to Bearded Vultures remains untested. Measuring their reflectance spectra would be necessary to determine how their chromatic properties relate to the spectral sensitivity and discrimination abilities of Bearded Vultures.

Overall, our findings illustrate that identifying the sensory mechanisms underlying bird collisions can fundamentally inform how mitigation measures are designed and emphasize that no universal visual solution is likely to reduce collisions across all species and environments. In Bearded Vultures, collision risk is unlikely to arise primarily from a lack of frontal vision, as the binocular field encompasses the forward flight path, even when the head is pitched downward during foraging. This contrasts with *Gyps* vultures (Martin et al. 2012) and Red Kites (Potier et al. in revision), in which a downward head orientation can place the forward flight path within the blind area. Likewise, their large eyes suggest relatively high spatial resolving power, which does not appear to be a limiting factor in the perception of aerial cables or visual mitigation markers equipped on cables. Instead, the particularly low contrast sensitivity of the Bearded Vulture may represent the most important sensory constraint, considerably reducing the detectability of cables and other aerial structures under natural viewing conditions. At the same time, the broad spectral sensitivity of its cone visual pigments suggests that mitigation devices could exploit both chromatic signals extending into the yellow-orange range and achromatic signals extending into longer, red wavelengths. Together, these results suggest that visibility rather than visual-field coverage or spatial resolution may represent the more important sensory limitation for obstacle detection in this species.

Taken together, our study supports the value of characterizing species-specific visual specializations and constraints when developing conservation strategies. Integrating complementary aspects of sensory ecology can help guide the design and experimental testing of mitigation devices more closely to the perceptual abilities of the species at risk (see Table S4).

## Limitations of the study

This study provides a first characterization of the visual capabilities of a protected and difficult-to-access species. The limited number of studied individuals precludes assessment of inter-individual variation and warrants caution when generalizing some of these measurements to the species as a whole, especially considering its wide and disjunct distribution across the Palearctic, Afrotropical and Indomalayan realms (Figure 1). Although only three individuals were examined, and contrast sensitivity could be fully assessed in only one individual, such sample sizes are common in studies of avian visual ecology (Reymond 1985; Potier et al. 2018a; Blary et al. 2024) and visual field measurements have been shown to be highly consistent both within and among individuals, indicating that relatively small sample sizes can provide reliable estimates of species-level visual characteristics (Lucas and Portugal, 2025).

The spectral limits reported here should be interpreted as theoretical estimates rather than absolute behavioural limits of vision. The potential cone-detection and chromatic-discrimination ranges were inferred from visual-pigment absorbance using a 5% α-band sensitivity criterion and do not incorporate ocular-media transmission, oil-droplet filtering, relative photoreceptor abundance and receptor noise. In particular, the proposed chromatic-discrimination range indicates wavelengths over which overlapping sensitivity of at least two cone pigments could support colour discrimination, rather than demonstrating behavioural colour discrimination at these spectral limits. Direct measurements of ocular transmission and oil-droplet absorbance, together with behavioural discrimination experiments, would be required to establish the functional limits of colour vision in Bearded Vultures.

## Data statement

All data are available in the electronic supplementary materials or at Figshare.

## Acknowledgements

The authors thank Johannès Thomas, Carmen Calero and Martín Chivite for their help with the fieldwork, Pierre Bouffandeau, Catherine Desnos and Pascale Malchiodi for logistical and technical assistance, and the National Eye Institute for providing 11,*cis*-retinal. Computing analyses were run on the GIGA cluster at ULiège. The study was funded by the Vanoise National Parc as part of a European project entitled POIA BirdSki, with the assistance of the European Union and the National Agency for Cohesion of Territory under the Interregional Convention of the Alps massif and in partnership with ASTERS-CEN74 in connection with a financial contribution of the Poma Foundation, with support from the FNADT – Alps Massif (National Fund for Territorial Planning and Development), and grants from the Belgian F.R.S.- FNRS (MISU F.6002.22 to JML and MISU F.6002.24 to MAL).

## Notes

### Competing Interest Statement

The authors have declared no competing interest.

