## Supplementary Information for "Seeing danger? Visual constraints and collision risk in the Bearded Vulture *Gypaetus barbatus*"

This file contains:

Supplementary methods

Supplementary Tables S1-S4

Supplementary Figures S1-S3

Supplementary data 1: opsin sequences

Supplementary references

### Supplementary methods

#### Contrast sensitivity

*Experimental set-up*

We used a reflex-like behaviour setup to study the contrast sensitivity of each individual, following Wagner et al. (2022) and Blary et al. (2024). This approach yields peak contrast sensitivities comparable to those obtained using operant conditioning (Blary et al., 2024), although it does not provide the spatial frequency associated with peak sensitivity. The optocollic reflexes (OCRs or nystagmus) are caused by involuntary compensatory response for image stabilization, *i.e.,* the individual follows the moving stimulus to keep the image stationary on the retina (Wagner et al., 2022). Because eye movements are limited in most birds (between 2 and 18°: Martin, 2009, O'Rourke et al., 2010), they track the object with conspicuous head movements.

To estimate the OCR, birds were placed individually into a U-shaped experimental box adapted to the size of the species (100 x 50 x 150 cm; L x W x H), with a matte black wooden floor and ceiling, and 3-mm thick clear polycarbonate walls. The transparent walls allowed the bird to see the stimulus, without affecting visual performance (Blary et al., 2024). The ceiling consisted of two wooden panels separated by 5 cm: a perforated panel allowed ventilation, while a smaller solid panel prevented the bird from seeing through the ventilation holes. All wooden surfaces were painted matte black (Liberty black Matt, GoodHome). A wooden door placed at the rear of the box was used to position and remove the bird.

The experimental box was placed at the centre of an octagonal array of computer monitors with a radius of 69cm. The array consisted of 28-inch monitors (Acer KG1 Series, 62x34 cm, 4K 3840x2160 pixels, 60 Hz, Acer Inc., Taiwan) mounted in portrait orientation, with their centres placed level with the bird's head. The displayed stimulus covered 236º horizontally and 46º vertically of the visual field. This setup provided sufficient visual-field coverage to elicit an OCR (Schmid & Wildsoet, 1998, Blary et al., 2024).

The monitor array and experimental box housing the bird were placed in an opaque black cloth tent (300 x 300 x 250 cm; L x W x H) with the base of array and the floor also covered by an opaque black cloth to minimize possible disturbance by external visual cues. The experimenter remained outside the tent throughout the experiment. The bird's behaviour was recorded from above using a video camera at 25 frames per second (Sony FDR-AX43 4K). In addition, a smartphone (Iphone 13, Apple Inc., USA) connected to a display outside the tent provided a live video feed, allowing the experimenter to monitor the bird throughout the procedure.

*Contrast sensitivity : Stimuli*

Stimuli were generated in Matlab (R2021) using Psychtoolbox (Brainard, 1997) and displayed simultaneously on the monitors using an 8-port HDMI splitter (4K, 60Hz, ST128HD20, StarTech). Stimuli comprised vertical sinusoidal achromatic gratings with varying spatial frequency and Michelson contrast. Gratings moved at an angular velocity of 20 degrees per second, similarly to values used for closely related species (Blary et al., 2024). To characterize contrast sensitivity, we tested six spatial frequencies (0.59, 0.85, 1.06, 1.27, 1.59 and 2.12 cycles per degree) over a range of Michelson contrasts from 0.012 and 1 (Table S2). Specifically, Michelson contrast ($Cm$) was defined as:

$$Cm = \frac{I_{max} - I_{min}}{I_{max} + I_{min}}$$

where $I_{max}$ and $I_{min}$ represent the maximum and minimum luminance of the grating, respectively. For each contrast level, the contrast was evaluated by measuring the difference in luminance between the darkest and lightest area of the grating at the centre of the monitor using a luxmeter (Hagner ScreenMaster, B. Hagner, Sweden) and expressed in candelas per square meter (cd/m$^2$).

Each stimulus was presented for 20 seconds (Schmid & Wildsoet, 1998, Shi & Stell, 2013, Wagner et al., 2022, Blary et al., 2024). Stimulus direction (clockwise or anticlockwise) was varied in a pseudorandom order, with no more than three consecutive trials in the same direction. illuminance measured at the bird's head was 110 lux inside the U-shaped box and 320 lux outside the box. Mean stimulus brightness was 73 cd/m$^2$.

*Contrast sensitivity : Experimental procedure*

Monitors were switched on 20 minutes before the beginning of each experiment to allow their luminance to stabilize. After being placed in the box, each bird was allowed to habituate for 10 min before the testing session began. The experimenter visually assessed the presence or absence of OCR following standard procedures (Schmid & Wildsoet, 1998, Shi & Stell, 2013, Blary et al., 2024). Each session started with maximum contrast (*i.e.* Michelson contrast = 1) to confirm that the bird exhibited a readily detectable OCR. Contrast was subsequently decreased and increased stepwise, following the procedure described by Blary *et al.* (2024), to determine the contrast threshold at which birds no longer exhibited the OCR. Each trial consisted of a specific combination of spatial frequency and contrast and was separated from the subsequent trial by an interval of at least 5 seconds during which an isoluminant homogeneous grey field was displayed. We used a four-of-five criterion (Shi & Stell, 2013, Blary et al., 2024). Basically, each frequency × contrast combination was presented five times, and considered reliably detected when an OCR was observed in at least four out of the five trials. The lowest contrast satisfying this criterion was defined as the threshold contrast for that spatial frequency. Once the threshold was determined, the procedure was repeated at a different spatial frequency while maintaining the same angular velocity. All sessions were video recorded, allowing the experimenter to review retrospectively the trials to confirm the presence or absence of OCR before analysis.

#### Prediction of opsin protein sequences

Opsin protein sequences were predicted from the publicly available Bearded Vulture genome assembly bGypBar2.pri (NCBI accession GCA_028022735.1). Gene models were predicted using the protein-profile extension of AUGUSTUS (AUGUSTUS-PPX; Keller et al., 2011), which incorporates information from alignments of homologous proteins to guide gene prediction. Reference avian protein sequences representing the major opsin families were retrieved from NCBI's Protein database and assembled into separate datasets, including rhodopsin, red-sensitive opsin, short-wave-sensitive opsin, melanopsin, pinopsin, opsin 3, opsin 5 and vertebrate ancient opsin. Sequences of each opsin family were aligned using MAFFT v7.526 (Katoh et al. 2002), and the resulting multiple-sequence alignments were processed to generate family-specific protein profiles for AUGUSTUS-PPX. Protein block profiles were generated using the AUGUSTUS *msa2prfl.pl* utility, with a maximum block-column entropy of 0.75 (*--max_entropy=0.75*).

To identify candidate genomic regions containing opsin genes, the reference avian opsin protein sequences were combined and aligned against the *G. barbatus* genome using BLAT (Kent 2002) in translated protein-to-genome mode *(-q=prot -t=dnax*). Overlapping BLAT hits were merged using BEDTools (Quinlan & Hall, 2010) to define candidate genomic intervals, which were subsequently extracted from the genome assembly and analysed with AUGUSTUS-PPX. Gene prediction used the chicken AUGUSTUS species model and the family-specific protein profiles, with protein-model thresholds set to *block_threshold_spec = 4.0*, *block_threshold_sens = 0.4* and *blockpart_threshold_spec = 4.5*; non-canonical GC donor splice sites were permitted. Predicted coding sequences and their translated protein products were recovered from the resulting AUGUSTUS GFF3 output for subsequent identification and analysis.

#### Visual opsin characterization

*Immunofluorescence validation of opsin expression and localization*

HEK293T cells were seeded onto coverslips in 6-well plates at a density of $0.6 \times 10^6$ cells per well containing Dulbecco's Modified Eagle Medium (DMEM, high glucose; Biowest) supplemented with 10% fetal bovine serum (FBS, Biowest). After 24h, at approximately 80% confluency, cells were transfected using polyethylenimine (PEI MAX, 40,000 MW; Polysciences) at a DNA:PEI ratio of 1:2. Specifically, 3 µg plasmid DNA and 6 µL PEI (1 mg/mL in water) were diluted in warm Opti-MEM reduced serum medium (Gibco). At 48 h post-transfection, coverslips were collected and washed three times with 1× D-PBS (Gibco), followed by fixation with 1 mL 4% paraformaldehyde (ThermoFisher) for 15 min at room temperature. Cells were rinsed three times with D-PBS for 5 min each and permeabilized with 0.25% Triton X-100 (Invitrogen) for 5 min at room temperature. Following three additional washes with D-PBS, a hydrophobic barrier was drawn around each coverslip using a PAP pen (Ted Pella 22311). Cells were blocked with 100 µL 10% goat serum (Life Technologies) for 30 min at 37 °C. The blocking solution was removed, and cells were incubated with an anti-Flag primary antibody (catalog number F1804, Merck) (1:50 dilution in D-PBS) overnight at 4°C or for 2 h at 37°C. Following three washes with PBST (D-PBS containing 0.1% Tween-20; Sigma-Aldrich), cells were incubated with Alexa Fluor 488 goat anti-mouse IgG (H+L) secondary antibody (1:200 dilution in D-PBS (catalog number A11001, Invitrogen) for 1 h at room temperature. Cells were then washed three times with D-PBS. Nuclei were counterstained with DAPI (5ug/ml) (ThermoFisher) for 5 min in the dark. After four washes with D-PBS, coverslips were mounted onto microscope slides using 5 µL antifade mounting medium (MCE).

Fluorescence images were acquired using an Echo Revolve fluorescent microscope (Echo). Immunofluorescence was used as a qualitative quality-control step to verify recombinant opsin expression and localization at the plasma membrane prior to visual-pigment characterization (Figure S3).

*Heterologous expression in HEK293T cells*

The open reading frames of the identified visual cone opsins (SWS1, SWS2, Rh2, LW) were codon-optimized, synthesized, and cloned into pUC57 by Genscript, with flanking *HindIII* and *BsiWI* restriction sites. After double restriction digestion with *HindIII* and *BsiWI,* the resulting inserts were separated by agarose gel electrophoresis and fragments of the expected sizes were gel-purified using the Monarch DNA gel extraction kit (NEB). Purified inserts were subcloned into the linearized pcDNA5-FLAG-T2A-mRuby2 expression vector (Liénard et al., 2021) and transformed into *Escherichia coli* 5-alpha competent cells (NEB) for plasmid propagation. Individual bacterial colonies were cultured in Luria-Broth (LB) medium containing ampicillin, plasmid DNA was purified, and the sequence of each construct was verified by Sanger sequencing. Plasmid stocks (480 µg per construct) were subsequently prepared following the ZymoPURE II Plasmid MidiPrep kit protocol (Zymo Research).

Rhodopsin complexes were reconstituted in human embryonic kidney cells (HEK293T) in presence of 11-*cis*-retinal following the PASHE procedure (Liénard et al., 2022). Absorbance spectra of purified rhodopsin were measured in the dark from 1.5-µL aliquots using an Implen UV-VIS spectrophotometer (ThermoFisher). Protein purification fractions were analysed by Western blot as described previously (Roberts et al., 2025) (Figure S2). For each rhodopsin, the wavelength of maximum absorbance ($\lambda_{max}$) was estimated by nonlinear fitting of the absorbance data to a visual pigment template (Govardovskii et al., 2000) in R v. 3.6.6. using the rsample (Frick et al., 2022) and tidymodels (Kuhn & Wickham, 2020) packages.

#### Supplementary tables

**Table S1:** Visual field data.

**Female Puy du Fou**

| Elevation (deg) | Binocular overlap (deg) |
|---|---|
| 350 | -0.9 |
| 0 | 16.1 |
| 10 | 14.1 |
| 20 | 13.1 |
| 30 | 19.0 |
| 40 | 23.0 |
| 50 | 24.0 |
| 60 | 34.9 |
| 70 | 38.9 |
| 80 | 33.9 |
| 90 | 30.9 |
| 100 | 30.0 |
| 110 | 21.0 |
| 120 | 11.1 |
| 130 | -8.0 |

**Male Vallcalent**

| Elevation (deg) | Binocular overlap (deg) |
|---|---|
| 10 | 1.7 |
| 20 | 9.7 |
| 30 | 9.7 |
| 40 | 21.7 |
| 50 | 23.6 |
| 60 | 27.6 |
| 70 | 28.6 |
| 80 | 38.5 |
| 90 | 26.6 |
| 100 | 17.7 |
| 110 | 16.7 |
| 120 | 19.7 |
| 130 | 3.7 |

**Female Vallcalent**

| Elevation (deg) | Binocular overlap (deg) |
|---|---|
| 0 | -3.0 |
| 10 | 7.8 |
| 20 | 15.8 |

| | |
|---|---|
| 30 | 22.7 |
| 40 | 24.7 |
| 50 | 29.7 |
| 60 | 28.7 |
| 70 | 34.6 |
| 80 | 26.7 |
| 90 | 23.7 |
| 100 | 21.7 |
| 110 | 19.7 |
| 120 | 3.8 |

166

**Table S2:** Steps used for the contrast sensitivity estimation and their associated contrasts (of the vertical sinusoidal achromatic gratings) and the average luminance of monitors.

| Step | Michelson Contrast | Inverse Michelson Contrast | Average luminance |
|---|---|---|---|
| 127 | 1.00 | 1.00 | 130.26 |
| 120 | 1.00 | 1.00 | 124.79 |
| 110 | 0.99 | 1.01 | 116.42 |
| 100 | 0.97 | 1.03 | 107.09 |
| 90 | 0.94 | 1.07 | 99.86 |
| 80 | 0.90 | 1.11 | 93.51 |
| 70 | 0.85 | 1.18 | 87.80 |
| 60 | 0.78 | 1.29 | 82.73 |
| 50 | 0.69 | 1.45 | 77.88 |
| 45 | 0.64 | 1.57 | 75.36 |
| 40 | 0.58 | 1.71 | 73.18 |
| 35 | 0.52 | 1.92 | 71.24 |
| 30 | 0.46 | 2.17 | 69.79 |
| 29 | 0.45 | 2.24 | 69.33 |
| 28 | 0.43 | 2.31 | 68.97 |
| 27 | 0.42 | 2.39 | 68.53 |
| 26 | 0.40 | 2.47 | 68.28 |
| 25 | 0.39 | 2.55 | 67.95 |
| 24 | 0.38 | 2.65 | 67.89 |
| 23 | 0.37 | 2.74 | 67.31 |
| 22 | 0.35 | 2.88 | 67.21 |
| 21 | 0.33 | 2.99 | 67.06 |
| 20 | 0.32 | 3.13 | 66.67 |
| 19 | 0.30 | 3.29 | 66.57 |
| 18 | 0.29 | 3.45 | 66.34 |
| 17 | 0.27 | 3.64 | 66.09 |
| 16 | 0.26 | 3.84 | 65.88 |
| 15 | 0.24 | 4.10 | 65.57 |
| 14 | 0.23 | 4.39 | 65.33 |
| 13 | 0.21 | 4.71 | 65.21 |
| 12 | 0.20 | 5.07 | 64.98 |
| 11 | 0.18 | 5.50 | 64.71 |
| 10 | 0.17 | 6.04 | 64.62 |
| 9 | 0.15 | 6.62 | 64.49 |
| 8 | 0.13 | 7.50 | 64.13 |
| 7 | 0.12 | 8.51 | 63.96 |
| 6 | 0.10 | 9.62 | 63.78 |
| 5 | 0.09 | 11.46 | 63.57 |
| 4 | 0.07 | 13.89 | 63.50 |
| 3.5 | 0.06 | 16.12 | 64.05 |
| 3 | 0.05 | 18.32 | 63.54 |

| | | | |
|---|---|---|---|
| 2.5 | 0.05 | 21.70 | 64.04 |
| 2 | 0.04 | 26.73 | 63.35 |
| 1.5 | 0.03 | 35.39 | 63.89 |
| 1 | 0.02 | 51.29 | 63.23 |
| 0.5 | 0.01 | 82.54 | 63.76 |

**Table S3**: Contrast sensitivity data.

| Spatial Frequency (c/deg) | Inverse Cm Male Vallcalent | Inverse Cm Female Vallcalent | Average |
|---|---|---|---|
| 2.11 | 5.55 | | 5.55 |
| 1.59 | 7.14 | | 7.14 |
| 1.27 | 9.09 | 10 | 9.55 |
| 1.06 | 7.69 | | 7.69 |
| 0.84 | 4.54 | 5.88 | 5.21 |
| 0.59 | 6.25 | 5.76 | 6.01 |

**Table S4. Prediction of detection distances and visual marker size (following Martin 2022 principles).**

| Parameter | Value |
|---|---|
| ***Minimal distance to resolve two objects in flight ($D = 2\ s \times V$)*** | |
| Flight speed, V (Rushworth and Krüger 2014; Fluhr et al. 2025) | 11.1 m/s |
| Minimal distance, $D = 2\ s \times V$ | 22.2 m |
| ***Maximal estimated spacing between two adjacent objects, $d = 2D \cdot \tan(\theta/2)$*** | |
| Binocular overlap, $\theta$ (27° × π/180) | 0.471 rad |
| Half binocular overlap, $\theta/2$ (13.5° × π/180) | 0.236 rad |
| $\tan(\theta/2)$ | 0.240 |
| Maximal estimated spacing, d | 10.7 m |
| ***Minimal width (w) of an object, $w = D \cdot \tan(1/(2 \cdot SR))$*** | |
| 1/(2 × SR), with SR = 98.5 cycles/degree (see Table 1) | $5.08E^{-03}$ deg |
| π/180 (conversion from degrees to radians) | 0.0175 rad/deg |
| Minimal object width under good visibility | 0.20 cm |
| Minimal object width under poor visibility* | 20 mm |
| Achromatic 3 × 3 square pattern (see Martin 2022), minimal width (5.4 × 5.4 cm) | 60 mm |

*Note: visual resolution during twilight can be reduced by up to 10-fold compared with maximum spatial resolution (visual acuity) (Martin 2022). An animal would therefore have to be 10 times closer to detect a given object at twilight than at midday, or the object would have to be 10 times larger

### Supplementary Figures

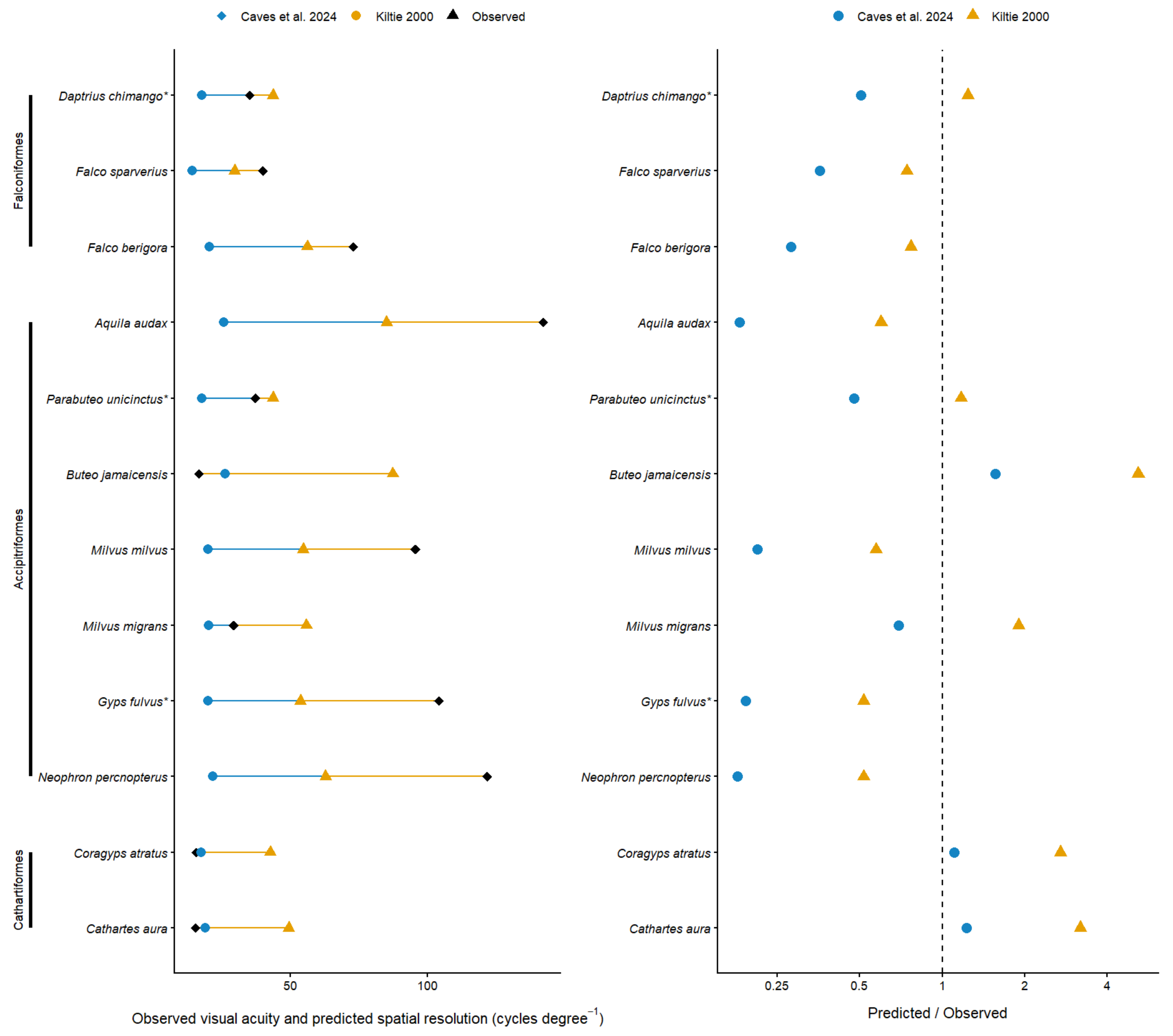


**Figure S1. Comparison of observed visual acuity and eye-size-based predictions of spatial resolution in diurnal raptors. *(left)*** Observed visual acuity and spatial resolution predicted from axial eye length (AL; mm) for diurnal raptor species included in the dataset compiled by Caves et al. (2024), supplemented with species for which AL was estimated from our own measurements of corneal diameter (*). Among these, measurements for *Parabuteo unicinctus* (Potier et al. 2016a) and *Daptrius chimango* (Potier et al. 2016b) have been previously published. Predictions were calculated using the relationships suitable for diurnal birds from Caves et al. (2024) and Kiltie (2000). Black diamonds indicate observed visual acuity, blue circles and orange triangles predictions-based Caves et al. (2024) and Kiltie (2000), respectively. **(*Right*)** Ratio of predicted spatial resolution to observed visual acuity for each species. The vertical dashed line indicates a ratio of 1, corresponding to perfect agreement between predicted and observed values; values below and above 1 indicate under- and over-estimation, respectively. Note that the x-axis is logarithmically scaled.

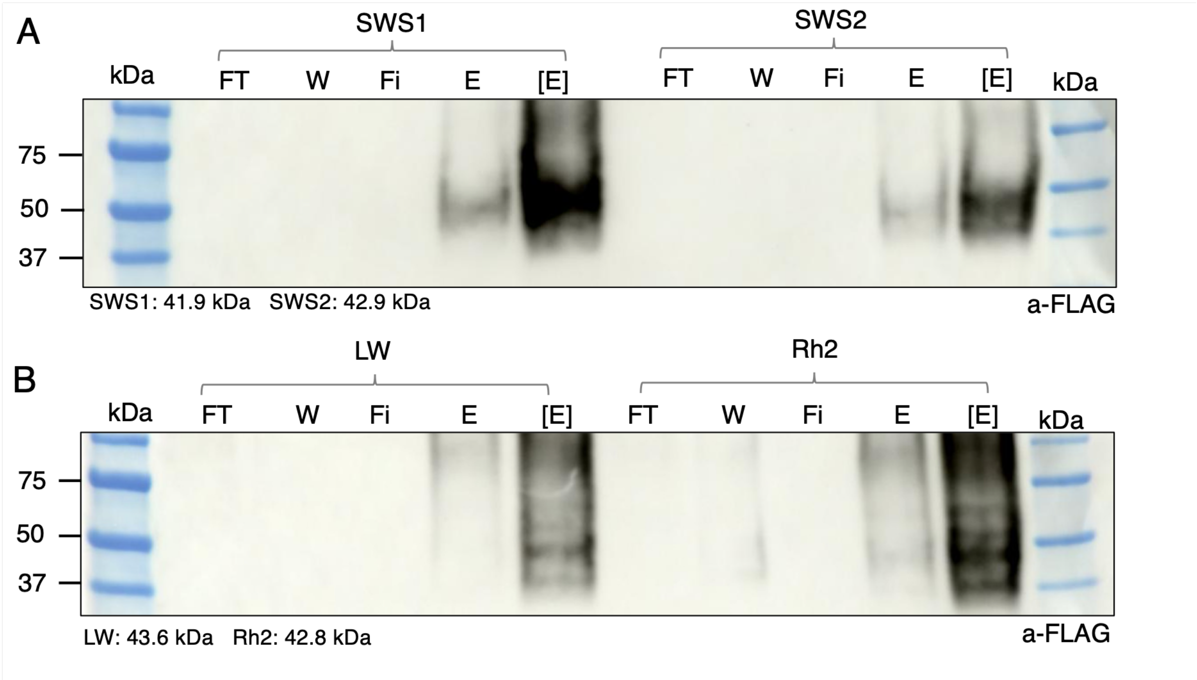


**Figure S2**. **Western blot analyses of opsin purification fractions.** (**A**) SWS1 and SWS2; (**B**) LW and Rh2. FT, flow through containing cellular proteins not retained during opsin purification; W, wash fraction to remove residual unbound retinal; Fi, filtrate from the concentration column; E, unconcentrated FLAG protein eluate; [E], concentrated opsin fraction. Expected molecular masses are indicated below each blot.

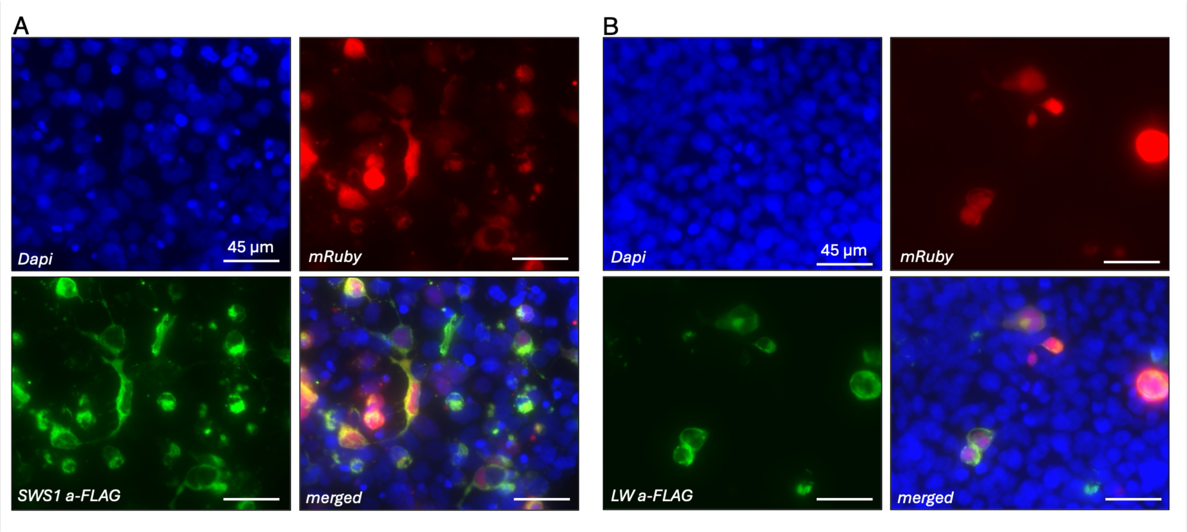


**Figure S3**. **Immunofluorescence validation of recombinant opsin expression and localization in HEK293T cells** Representative fluorescence images of fixed HEK293 cells 48 h post-transfection with constructs encoding. (**A**) SWS1 or (**B**) LW opsins. Recombinant opsin proteins were detected using an anti-FLAG antibody (green), showing predominantly plasma-membrane-associated localization. mRuby2, which is expressed from the same construct, served as an intracellular reported (red). Nuclei were counterstained with Dapi (blue). Scale bars: 45 µm.

#### Supplementary Data 1:

*Predicted opsin amino acid sequences*:

>Gba_SWS1 (CM051535.1:1320427-1323762|short-wave-sensitive|g25; chromosome 4)
MSGDEEFYLFKNGSSVGPWDGPQYHIAPPWAFYLQTAFMGFVFLVGTPLNAIVLVVTVKYKKLR
QPLNYILVNISFSGFISCIFSVFTVFVSSSQGYFVFGKHMCSLEGFVGATGGLVTGWSLAFLAFERYI
VICKPFGNFRFSSKHALMVVVATWVIGIGVAIPPFFGWSRYMPEGLQCSCGPDWYTVGTKYKSEY
YTWFLFIFCFIVPLSLIIFSYSQLLSALRAVAAQQQESATTQKAEREVSRMVVVMVGSFCLCYVPYA
ALAMYMVNNRNHGLDLRLVTIPAFFSKSACVYNPIIYCFMNKQFRACIMETVCGKPMTDDSDISSS
AQRTEVSSVSSSQVSPS
>Gba_SWS2 (CM051559.1:19629-23583|short-wave-sensitive|g3; chromosome 28)
MQKSREGREELPEDFYIPMALDTPNLTALSPFLVPQTHLGSPGVFMGMSAFMFMLIALGVPINTLTI
FCTAKYKKLRSHLNYILVNLAVSNLLVICVGSTTAFYSFSQMYFALGPTACKIEGFAATLGGMVSL
WSLAVVAFERFLVICKPLGNFTFRGSHAVLGCAATWIFGLVASVPPLFGWSRYIPEGLQCSCGPDW
YTSNNKWNNESYVIFLFCFCFGVPLAIIIFSYGRLLLTLRAVAKQQEQSATTQKAEREVTKMVVVM
VLGFLVCWAPYSAFALWVVTHRGQAFDVGLASIPSVFSKASTVYNPVIYVFMNKQFRSCMLKLVF
CGKSPFGDEDDVSGSSQATQVSSVSSSQVAPA
>Gba_Rh2 (CM051552.1:4605531-4610331|rhodopsin|g47; chromosome 21)
MNGTEGINFYVPMSNKTGVVRSPFEYPQYYLAEPWKYRVVCCYIFFLISTGLPINLLTLLVTFKHK
KLRQPLNYILVNLAVADLFMACFGFTVTFYTAWNGYFVFGPVGCAVEGFFATLGGQVALWSLVV
LAIERYIVVCKPMGNFRFSATHAMMGIAFTWIMAFSCAAPPLFGWSRYMPEGMQCSCGPDYYTH
NPDYHNESYVLYMFVIHFIIPVVVIFFSYGRLICKVREAAAQQQESATTQKAEKEVTRMVILMVLG
FMLAWTPYAVVAFWIFTNKGADFTATLMSVPAFFSKSSSLYNPIIYVLMNKQFRNCMITTICCGKN
PFGDEDTSSAVSQSKTEVSSISSSQVSPA
>Gba_LW (CM051559.1:14013-16259|red-sensitive|g2; chromosome 28)
MAAGWEAAVFAARRRHEDEDTTRDSVFTYTNSNNTRGPFEGPNYHIAPRWVYNLTSLWMIFVVV
ASIFTNGLVLVATWKFKKLRHPLNWILVNLAVADLGETVIASTISVINQISGYFILGHPLCIIEGYTV
SACGITALWSLAIISWERWFVVCKPFGNIKFDGKLAVAGILFSWIWSCAWTAPPIFGWSRYWPHGL
KTSCGPDVFSGSSDPGVQSYMVVLMVTCCFFPLAIIILCYLQVWLAIRAVAAQQKESESTQKAEKE
VSRMVVVMIVAYCFCWGPYTIFACFAAANPGYAFHPLAAALPAYFAKSATIYNPIIYVFMNRQFR
NCILQLFGKKVDDGSEVSTSRTEVSSVSNSSVSPA
